# Historeceptomics Profiling of Ketamine

**DOI:** 10.64898/2026.11.18.767642

**Authors:** Eric Rosenn, Harrison Agnor, Jennifer Fuller, Timothy Cardozo

## Abstract

Ketamine elicits distinctive neuropsychiatric phenotypes from a complex and as yet elusive mechanism of action. Here, we applied an updated Historeceptomics approach to identify molecular targets and brain tissues that may contribute to its antidepressant, analgesic, and dissociative effects. Historeceptomics generates multidimensional target-tissue fingerprints by integrating drug-target activity with tissue-specific target expression, providing an anatomic profile of drug action. The results suggest that ketamine activity on NMDA receptors in the prefrontal and cingulate cortices and the amygdala may contribute to ketamine improvements in social functioning for MDD. However, this same receptor-tissue activity may invoke hallucinatory effects at higher intensity or with engagement of additional brain regions. The central nervous system (CNS) fingerprint for ketamine was compared with those of other psychoactive drugs, showing overlap with fingerprints from other antidepressant, hallucinogenic, and analgesic drugs. Some ketamine effects may be traced to specific enantiomers or metabolites of ketamine. These findings provide new insights into the mechanisms by which ketamine elicits its unique phenotype and demonstrate the utility of using Historeceptomics to understand complex drug action.

## INTRODUCTION

Ketamine was developed as the anesthetic CI-581 in the early 1960s and approved by the U.S. Food and Drug Administration for general anesthesia in 1970.^1,2^ Its use has since extended to pain management and depression, while its dissociative and perceptual effects have also contributed to recreational use and abuse.^3,4^ Initial randomized trials showed that a single intravenous ketamine infusion could produce improvement in clinical depression scores within hours, with effects lasting several days.^5,6^ Subsequent meta-analyses and real-world studies have supported its short-term antidepressant efficacy, although repeated or maintenance dosing is generally required to prolong responses.^7–9^ Intranasal esketamine, the S-enantiomer of ketamine, received FDA approval for treatment-resistant depression in 2019.^10^

Ketamine administration results in varied phenotypes (**Table S1**). Several components of ketamine’s mechanism of action (MoA) for these phenotypes have been suggested.^11,12^ At subanesthetic concentrations, inhibition of N-methyl-D-aspartate receptors (NMDARs) on GABAergic interneurons can reduce inhibitory input to pyramidal neurons, increasing glutamate release, enhancing AMPA receptor signaling, and promoting synaptic plasticity associated with antidepressant responses.^13–15^ NMDAR antagonism, however, does not fully account for ketamine’s pharmacology or rapid antidepressant effects.^16–18^ Ketamine also interacts with muscarinic and nicotinic acetylcholine receptors,^19,20^ opioid receptors,^21^ monoamine transporters,^22^ and dopamine and serotonin receptors.^23^ These additional activities, as well as the distribution of these receptors in specific brain tissues, may contribute to its antidepressant, analgesic, dissociative, and cognitive effects.^24–26^

Previously, we developed historeceptomics (HR), which aims to incorporate anatomy into drug MoA by integrating drug-target affinities with tissue-specific target expression to generate multidimensional target-tissue fingerprints.^27,28^ The method was applied to clozapine and bupropion, newly identifying some shared and drug-specific target-tissue relationships that may help explain some of their therapeutic and adverse effects.^29,30^

To further characterize ketamine’s MoA, we applied the HR approach to characterize the central nervous system (CNS) fingerprint of ketamine and its significant metabolites. We compared this fingerprint with those of selected psychoactive drugs and examined its receptor–tissue components in relation to published pharmacological, behavioral, and clinical studies.

## METHODS

### Drug bioactivity data

Quantitative ketamine-target activity data were obtained from ChEMBL,^31^ PubChem BioAssay,^32,33^ the NIMH Psychoactive Drug Screening Program Ki Database,^34^ the IUPHAR/BPS Guide to Pharmacology,^35^ BindingDB,^36^ and primary literature (**Table S2**).^37^ In addition to ketamine, bioactivity data were collected for racemic ketamine, S-ketamine, R-ketamine, hydroxyketamine, norketamine, (2R,6R)-hydroxynorketamine, and (2S,6S)-hydroxynorketamine. Targets were restricted to individual proteins mapped to standardized gene symbols. Activity measurements were converted to molar units and expressed as:

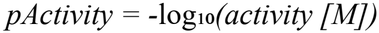

For each target, measured human data were used when available; otherwise, measured data from other mammalian species were used. Within the selected species, exact measurements were preferred to bounded measurements, and activity measurements were prioritized as Ki, Kd, IC50, EC50/AC50, followed by other quantitative potency measurements. When multiple eligible measurements remained, the strongest activity value was retained. Nonmammalian measurements and qualitative records were excluded from HR scoring.

### Tissue-specific gene expression data

Human tissue-specific gene expression data were obtained from the BioGPS GeneAtlas U133A/GNF1H GCRMA dataset.^38–40^ Probe identifiers were mapped to their corresponding genes, and ambiguous mappings were excluded. When multiple probes mapped to the same gene, the median expression value was used. Expression for each gene was standardized across the tissue panel as a Z-score. The CNS analysis was restricted to 18 nervous-system tissues represented in the BioGPS dataset.

### Target-tissue scores and statistical analysis

For each ketamine target and tissue, a historeceptomic score was calculated by combining drug-target activity with the standardized tissue expression of the corresponding target:

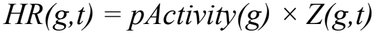

where pActivity(g) represents ketamine activity at target g, and Z(g,t) represents the standardized expression of that target in tissue t.^27^ Target-tissue pairs with unusually high HR scores were identified using the generalized extreme Studentized deviate test.^41^ The primary CNS fingerprint was defined at α = 0.001, and α = 0.0001 was examined as a more stringent threshold (**Table S3**).

### Drug fingerprint comparisons and principal component analysis

Ketamine was compared with selected psychoactive drugs representing antidepressant, antipsychotic, serotonergic psychedelic, anesthetic/sedative, dissociative, analgesic/opioid, mood-stabilizing, and cholinergic pharmacology. Fingerprint overlap (**Tables S6–S7** and **Figures S1–S3**) was assessed at α = 0.001 by identifying target–tissue pairs retained in both compound fingerprints. Shared components required matching gene symbols and tissue labels, irrespective of differences in HR score magnitude. Principal component analysis (PCA) was performed on a matrix of 35 compound profiles and 30 variable target–tissue features selected at α = 0.001. Features were encoded as binary fingerprint calls (1 = retained, 0 = observed but not retained), with unavailable target–tissue observations represented as missing values. PCA was performed using iterative expectation-maximization singular value decomposition (EM-SVD), with missing entries initialized from feature means and iteratively estimated while preserving observed values. The first two principal components were used to visualize relationships among compound fingerprints.^29,42^

## RESULTS

We combined available ketamine-target pharmacological measurements with human tissue-specific gene expression data to obtain target-tissue scores. An outlier-detection procedure identified the statistically significant scores defining ketamine’s CNS historeceptomic fingerprint (**Figure 1** and **Tables S4–S5**). NMDA- and GABA-A-receptor subunits were prominent components of the fingerprint, with distinct anatomical distributions. GRIN1 was represented in the prefrontal cortex and amygdala, while GABA-A-receptor features involved GABRA1 in the occipital and parietal lobes and GABRA2 in the amygdala. The prefrontal cortex also contained muscarinic CHRM3, serotonergic HTR2A, and phosphodiesterase PDE4A features. Additional target-tissue relationships extended to the hypothalamus, caudate nucleus, thalamus, and other CNS regions. Separate HR fingerprints for S-ketamine, R-ketamine, and hydroxyketamine were mapped to phenotypes (**Figure 2** and **Table S8**).

**Figure 1.**
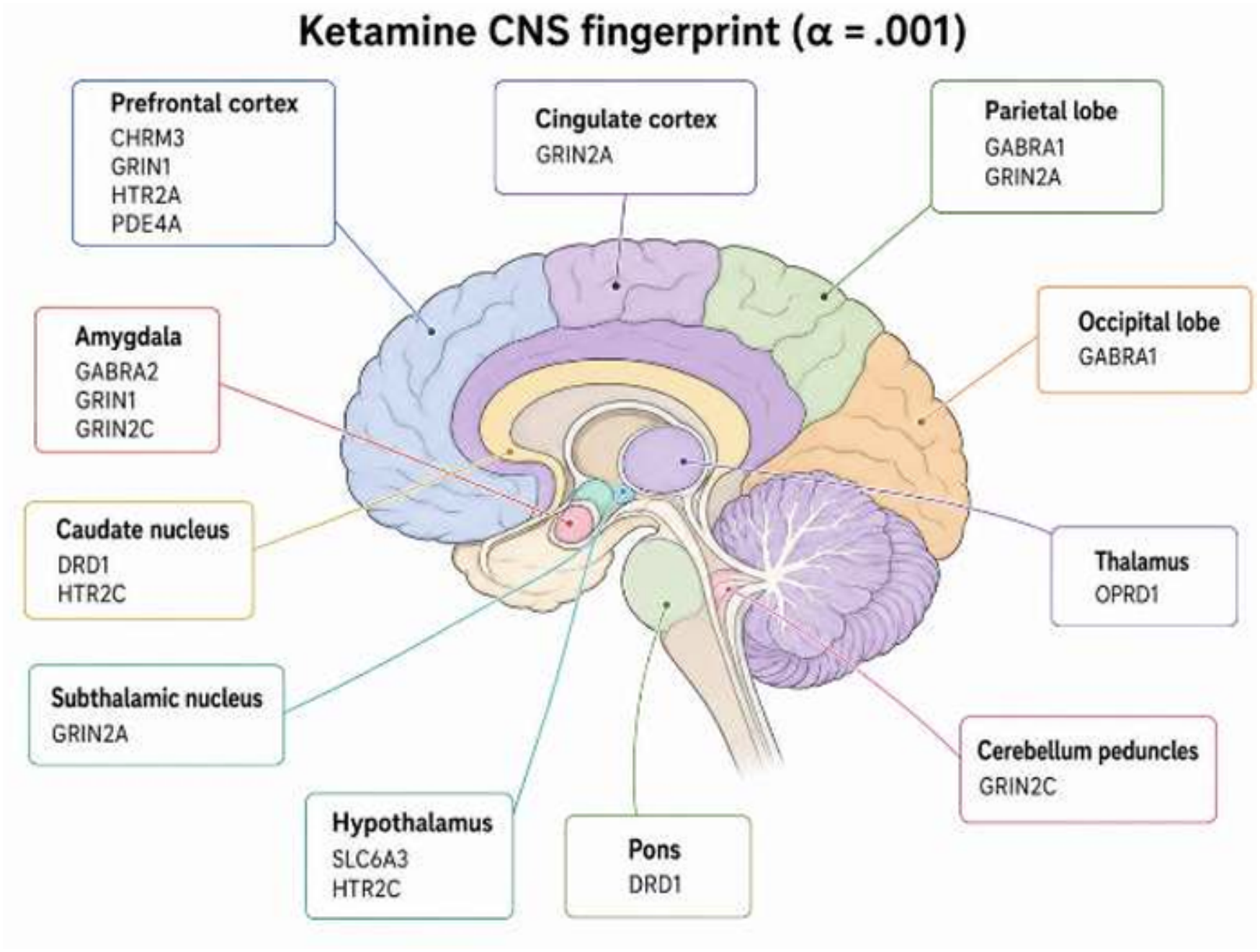
Ketamine central nervous system historeceptomic fingerprint. Sagittal brain schematic showing the 19 target–tissue components of the ketamine CNS fingerprint selected by the generalized extreme Studentized deviate test at α = 0.001. Labels identify the retained targets by gene symbol within each anatomical region. HR scores combine drug-target activity with standardized human tissue expression. GRIN1, GRIN2A and GRIN2C encode NMDA receptor subunits; GABRA1 and GABRA2 encode GABA-A receptor subunits. The remaining labels identify muscarinic, serotonin, dopamine and opioid receptors, the dopamine transporter SLC6A3, and phosphodiesterase PDE4A. Created with BioRender.

**Figure 2.**
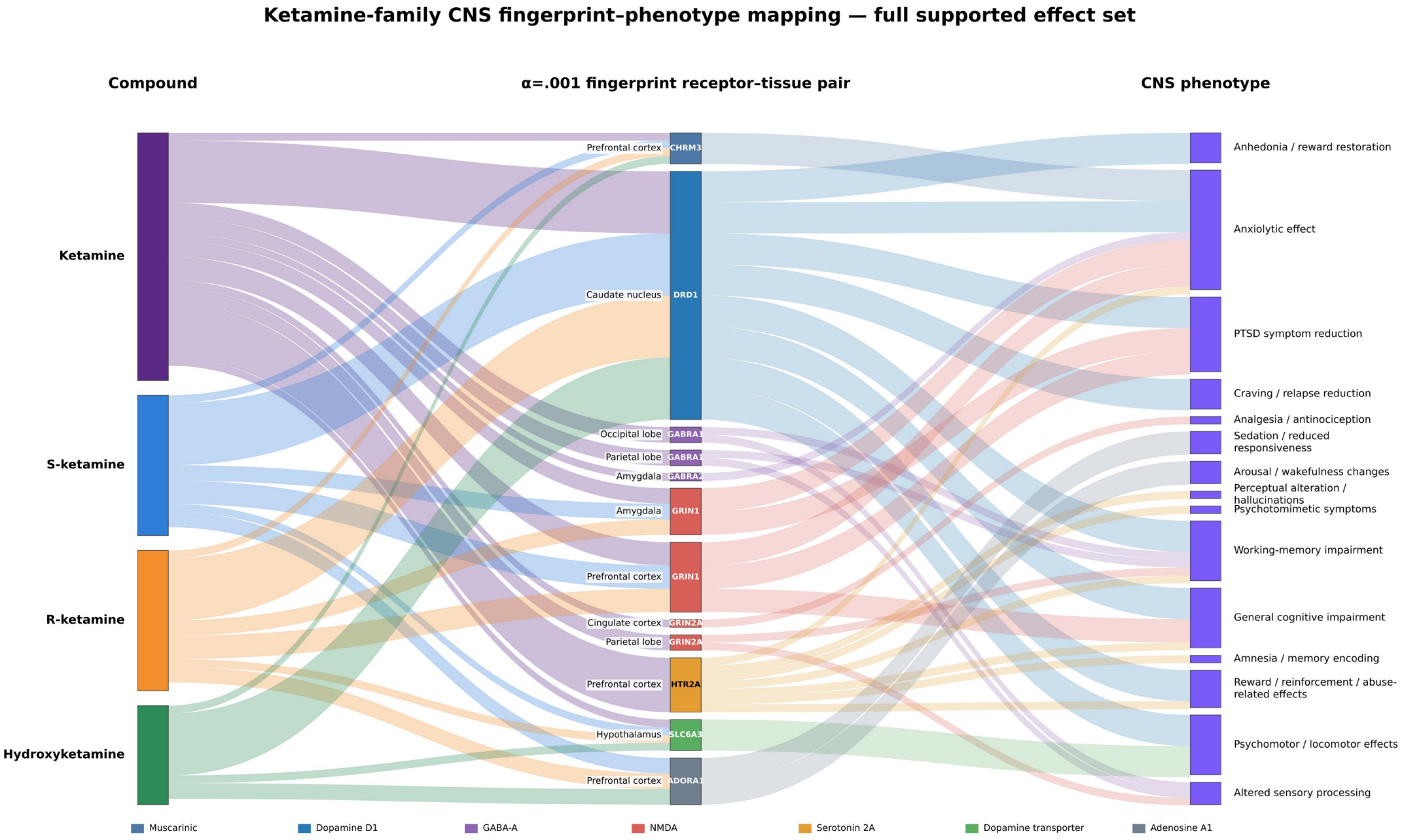
Full ketamine-family CNS fingerprint-phenotype mapping. Sankey diagram retaining all 15 exact phenotype categories and 76 compound-pair-phenotype paths. Ribbon width represents the number of represented paths and does not encode effect magnitude or evidence strength. This expanded figure preserves the exact phenotype categories underlying the compact phenotype mapping and documents the complete set of supported compound-pair-phenotype paths.

Comparison with psychoactive reference drugs identified shared fingerprint components with antidepressant, hallucinogenic, and analgesic compounds. CHRM3 in the prefrontal cortex was shared with fluoxetine, sertraline, and scopolamine, while the dopamine transporter SLC6A3 in the hypothalamus was shared with bupropion, fluoxetine, and sertraline. HTR2A in the prefrontal cortex was shared with LSD, psilocin, and psilocybin, and the delta-opioid receptor OPRD1 in the thalamus was shared with morphine. Several NMDA- and GABA-A-receptor pairs, together with PDE4A in the prefrontal cortex, were not represented in the reference-drug fingerprints. The comparisons therefore identified both shared receptor-tissue components and features that distinguished ketamine within the drugs examined.

The drug fingerprints were represented as target-tissue feature vectors and visualized by principal component analysis (**Figure 3**). Ketamine was positioned near bupropion, phencyclidine, and sertraline in the displayed projection. Cortical HTR2A and GABRA1 features contributed in opposite directions to the first principal component, while GRIN2C in the amygdala was the largest contributor by absolute loading to the second principal component.

**Figure 3.**
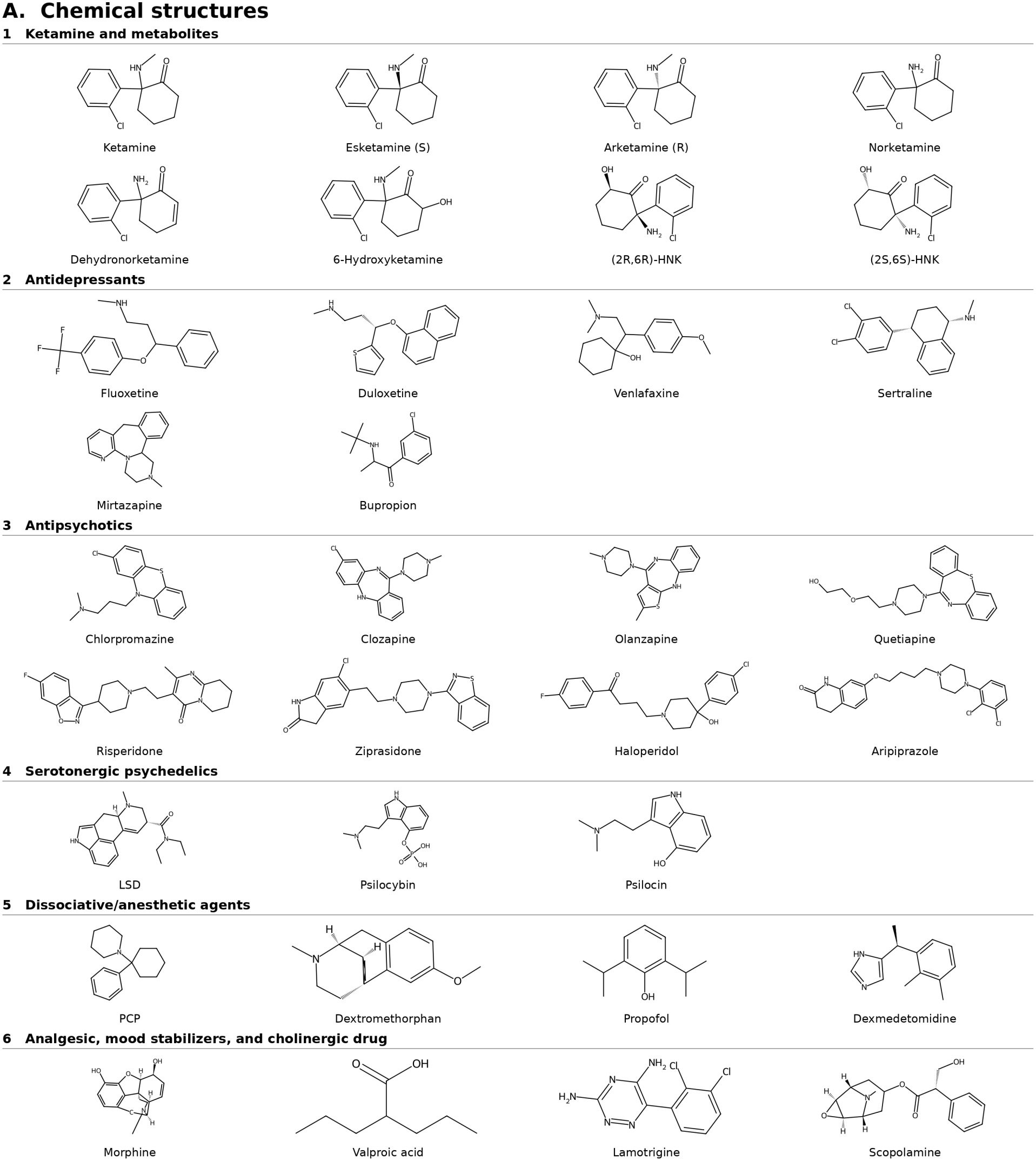
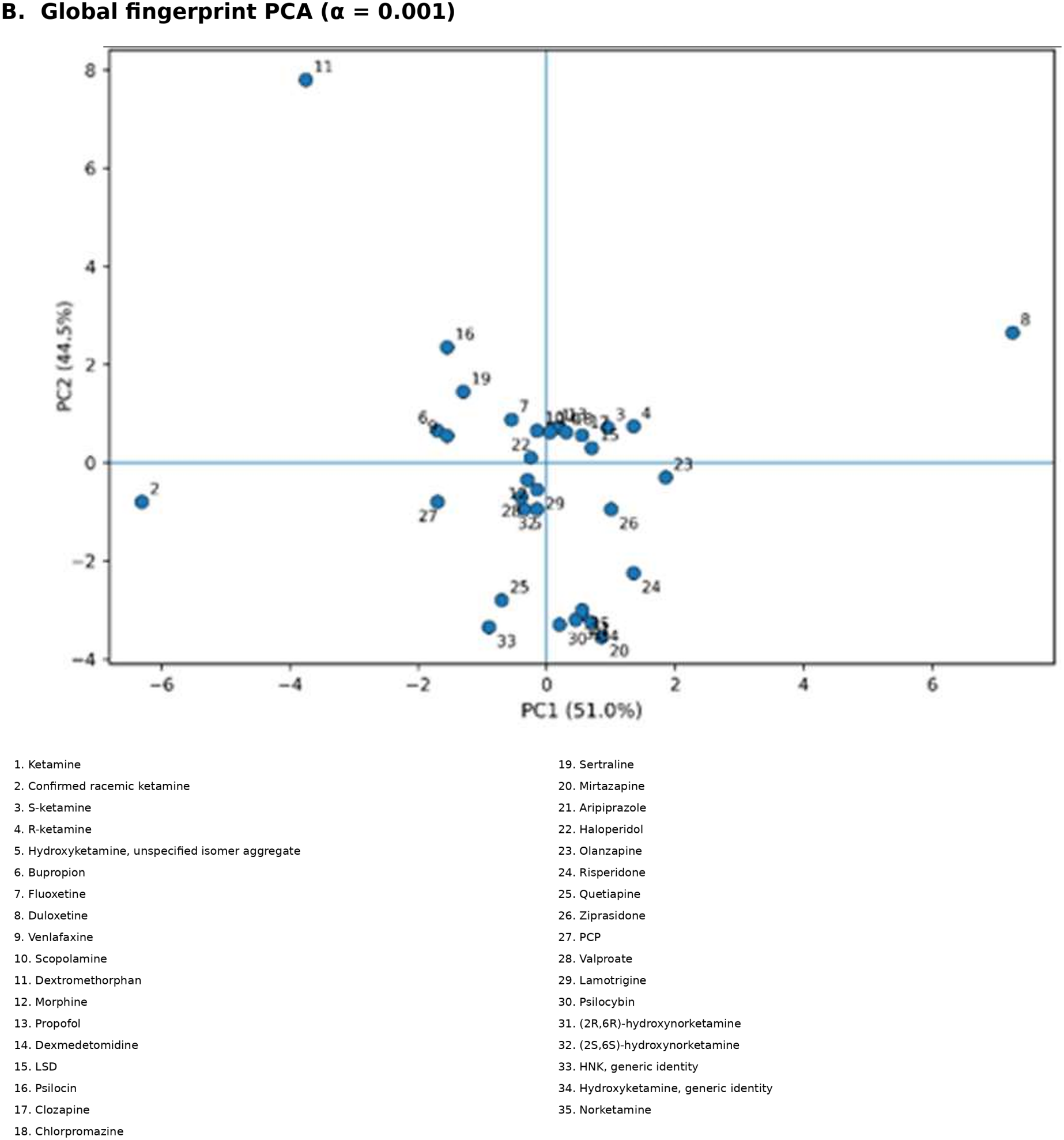
Chemical structures and fingerprint comparison of ketamine and psychoactive drugs. (A) PubChem-verified chemical structures of ketamine-related compounds and selected reference drugs, grouped by pharmacological category. The hydroxyketamine drawing represents the defined 6-hydroxyketamine positional isomer, not the unspecified-isomer aggregate. (B) Original principal component analysis of the global α = 0.001 fingerprint matrix, comprising 35 compound profiles and 30 variable target–tissue features. Numbered points correspond to the compound key. PC1 and PC2 account for 51.0% and 44.5% of the estimated variance, respectively.

## DISCUSSION

The ultimate goal of this report is to apply an updated HR approach^27,28^ to map the multiple affinities of ketamine for its receptors in brain tissues (**Figure 1**) to inform the MoA for some of its observable phenotypes. The finding that GRIN1 in the prefrontal cortex is a statistically dominant ketamine HR component is consistent with the well-established NMDAR involvement in ketamine’s antidepressant effects. Reducing GluN2B-containing NMDARs on medial-prefrontal GABA interneurons in mice blocked or occluded ketamine’s antidepressant-like effects.^43^ GRIN1 encodes the GluN1 subunit of NMDAR complexes, whose GluN2 subunit composition and cellular distribution influence receptor function.^44^ These receptor characteristics may help explain how prefrontal NMDAR inhibition could initiate antidepressant responses. Ketamine-induced prefrontal synapse formation and antidepressant-like behavior required mTOR activity.^45^ A brief change in transmission could therefore initiate a synaptic response that persists after acute cognitive effects have subsided.

Prefrontal, cingulate, and amygdalar NMDAR function may contribute to improvements in social functioning associated with ketamine treatment. A systematic review reported improvements in social functioning following ketamine or esketamine treatment.^46^ In mice, social defeat weakened functional connectivity between the medial prefrontal cortex and dorsal periaqueductal gray,^47^ while systemic ketamine restored social interaction and associated prefrontal neuronal activity in a subset of stress-susceptible animals.^48^ Selective *Grin1* deletion in anterior cingulate/prelimbic cortex increased preference for social novelty in mice without ketamine exposure.^49^ The parietal GRIN2A and GABRA1 components identified by HR suggest that excitatory and inhibitory signaling in the parietal cortex may modulate ketamine’s prefrontal antidepressant effects, consistent with reported changes in parietal-to-frontal AMPA-mediated connectivity and frontal GABA-A receptor dynamics.^50^ Human imaging studies associate antidepressant response to ketamine with prefrontal and cingulate function and connectivity,^51,52^ including normalization of reduced prefrontal connectivity and increased lateral-prefrontal and caudate connectivity in responders.^53^

The same brain regions that contribute to ketamine’s therapeutic social effects may also produce acute cognitive and perceptual disturbances. These can include working-memory impairment and hallucinations, with their severity varying according to dose and individual sensitivity.^26,54^ Within the cingulate, whose subregions contribute to pain and emotion processing,^55^ baseline subgenual perfusion was associated with next-day antidepressant improvement, whereas pregenual perfusion during infusion was associated with acute perceptual effects.^56^ Their different timing and anatomical distribution suggest that the processes associated with subsequent antidepressant improvement may be distinguishable from those producing acute perceptual disturbances. Ketamine also impaired working memory and reduced persistent firing in monkey dorsolateral prefrontal neurons.^57^ Inhibition of interneurons can increase pyramidal-cell activity, while interruption of recurrent excitation between pyramidal cells disrupts the sustained activity needed for working memory.^14,58^ Ketamine’s prefrontal GRIN1 fingerprint component may also be relevant to dose-dependent differences in cortical function, consistent with reports in rats that prefrontal glutamate release increases at subanesthetic doses but decreases at an anesthetic dose.^13^ The prefrontal 5-HT2A component shared with LSD, psilocin, and psilocybin may contribute to ketamine’s perceptual disturbances, potentially including hallucinations.^54,59,60^ Frontal 5-HT2A activity is prominent in serotonergic hallucinogen pharmacology,^59^ and its effects include changes in glutamatergic transmission. LSD and DOB inhibited NMDAR-mediated responses in rat prefrontal pyramidal neurons, and 5-HT2A antagonists blocked this effect.^61^ Serotonin-receptor activity can therefore alter NMDAR-mediated transmission within the same cortical neurons.^62^ Ketamine and serotonergic hallucinogens may consequently disturb prefrontal processing through different initial actions on interacting transmitter systems.^63^ The same prefrontal feature was also shared with clozapine and olanzapine, placing ketamine’s serotonergic pharmacology among compounds with contrasting psychoactive effects. This comparison is informative because drugs that produce different perceptual effects can act at the same receptor in the same region. Their functional effects at that receptor, together with their other targets, help explain why shared fingerprint components accompany different psychoactive profiles.

Prefrontal 5-HT2A activity likely also contributes to ketamine-induced short-term memory impairment. Local administration of the selective antagonist MDL100907 reversed this impairment in rats,^64^ identifying a way to modify cognition without directly reversing NMDAR blockade. However, 5-HT2A receptors also participate in psychedelic-induced plasticity,^65^ and deletion of medial-frontal receptors prevented dendritic spine formation and improvement in stress-related behavior after psilocybin.^66^ A similar contribution to ketamine-induced plasticity would mean that 5-HT2A blockade changes antidepressant activity as well as memory performance. For ketamine, newly formed prefrontal spines were required to maintain antidepressant effects on motivated escape behavior, although they were not required for the initial response.^67^ Separately, combined inhibition of the serotonin transporter and NMDARs produced rapid antidepressant-like effects involving prefrontal vasoactive intestinal peptide-expressing interneurons in mice.^18^

Thus, selectively preserving ketamine’s antidepressant and social-functioning effects while minimizing cognitive disturbances may require more precise targeting of NMDAR activity within specific cells and circuits of the prefrontal cortex, amygdala, and cingulate gyrus. Selected translational relationships between ketamine HR findings, psychiatric phenotypes, and potential development applications are summarized in **Table 1**.

**Table 1.**
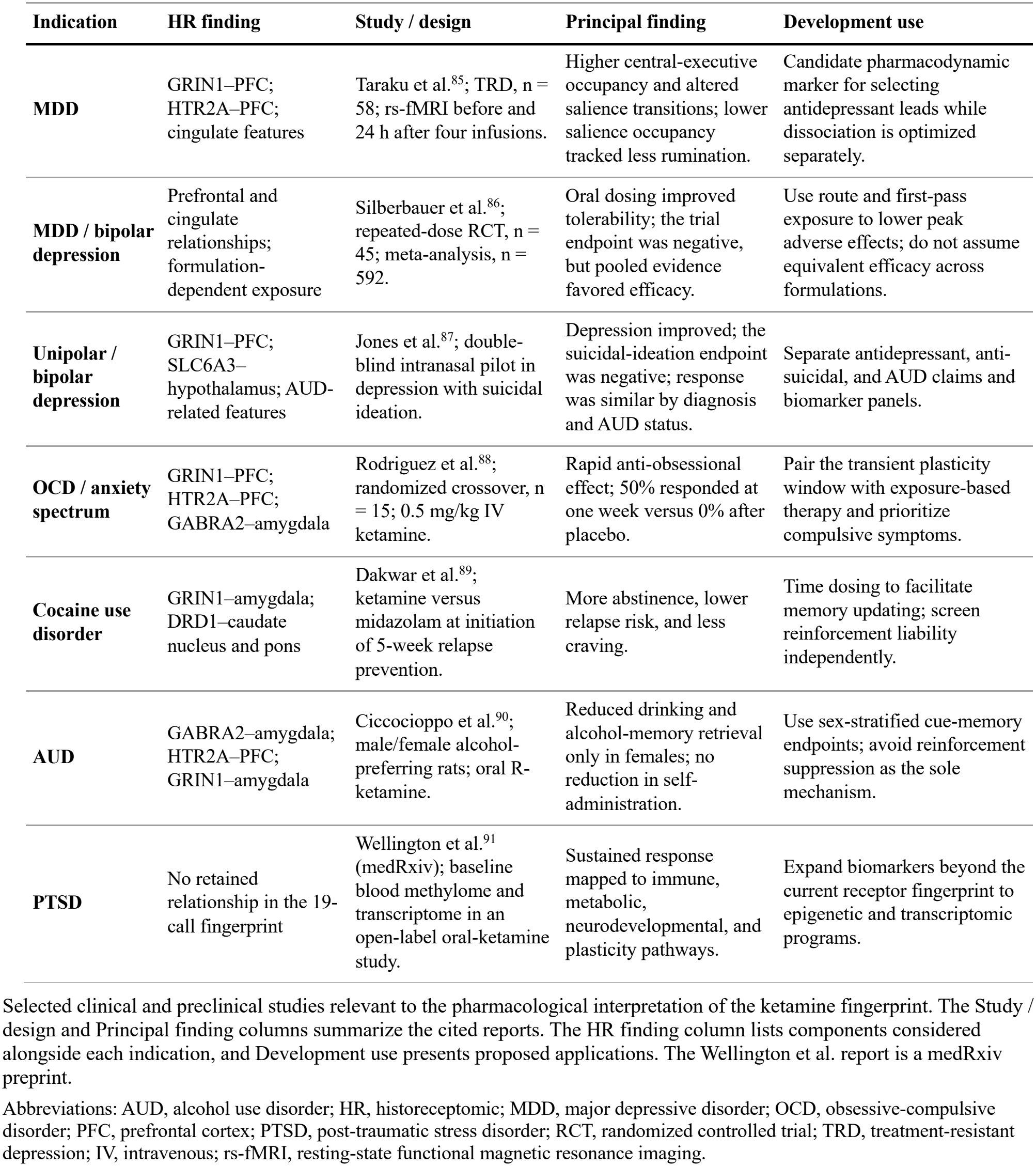
Selected translational relationships associated with the ketamine historeceptomic fingerprint.

Ketamine’s GRIN2C activity in cerebellar peduncles suggests a motor component of ketamine action that may be relevant to depression. GRIN2C-null mice had better motor coordination and markedly less ketamine-induced circling, while antidepressant-like responses were preserved. The proposed cerebellar involvement was based on receptor distribution and regional measurements; the experiments used global deletion and examined cerebellar cortex, not the peduncles specifically.^68^ GluN2A loss likewise produced antidepressant-like responses with limited psychotomimetic-like effects, with GluN2A-dependent changes in excitability demonstrated in hippocampal principal neurons.^69^ A compound with less activity at GluN2C-containing receptors might retain mood-related effects with less incoordination.

Ketamine has unique therapeutic activity in depression, including efficacy in treatment-resistant cases.^6,9,70–74^ Its rapid onset and potential effects on stress-related social behavior are important features of this therapeutic profile. Psychomotor retardation can occur in depression, including treatment-resistant cases.^75,76^ Ketamine’s HR fingerprint also includes prefrontal GRIN1, hypothalamic SLC6A3, and cerebellar-peduncle GRIN2C components that may together be relevant to the locomotor and other motor aspects of depression. Although ketamine has not been reported to improve psychomotor retardation consistently, the identified fingerprint components suggest that motor-related symptoms may warrant further examination in specific patient populations or clinical contexts.

Other findings of ketamine activity on specific targets in specific brain regions bear discussion, especially with respect to similarities with other psychiatric drugs. Hypothalamic dopamine transport, previously identified in bupropion and other antidepressant fingerprints,^30^ was shared with ketamine. Dopamine clearance is pharmacologically distinct from dopamine-receptor activity, and inhibition of monoamine transporters by ketamine has been demonstrated in cells expressing the human transporters.^22^ In mice, dopamine-transporter-expressing neurons in the hypothalamic periventricular nucleus link circadian input to locomotor control; inhibiting their dopamine output reduced amphetamine-induced hyperlocomotion.^77^ Altered dopamine handling in this region could therefore affect the behavioral activation accompanying mood improvement, including unwanted psychomotor activation.

Prefrontal muscarinic function may also contribute to ketamine’s antidepressant effects. Ketamine shared M3 targeting with scopolamine, fluoxetine, and sertraline despite differences in their principal pharmacology. Scopolamine produces rapid antidepressant effects in patients,^78^ and local administration into medial prefrontal cortex produces antidepressant-like behavior.^79^ Prefrontal GABA interneurons participate in this muscarinic response,^80^ providing a possible connection with ketamine-induced disinhibition. Ketamine’s antimuscarinic activity^20^ could reduce cholinergic excitation of inhibitory neurons and augment the response initiated by NMDAR blockade.

Muscarinic and glutamatergic actions would then jointly influence pyramidal-cell activity and subsequent plasticity. This offers a specific interpretation of the M3 overlap: cholinergic antagonism may modify the inhibitory control of prefrontal neurons even when the compounds differ in their other antidepressant mechanisms.

Thalamic δ-opioid activity may contribute to analgesia through the regulation of nociceptive transmission.^81,82^ Its overlap with morphine need not mean that ketamine acts as a conventional opioid agonist. Ketamine displaces ligands from recombinant opioid receptors, including δ receptors,^21^ and enhances responses to endogenous opioid peptides at μ, δ, and κ receptors in a receptor- and peptide-dependent manner.^83^ Enhancement of endogenous peptide activity offers an explanation for an opioid contribution despite weak direct agonist activity. In the thalamus, such modulation would depend on local peptide release as well as ketamine exposure, allowing endogenous pain control to act alongside reduced glutamatergic excitation. Opioid activity may also influence antidepressant responses. Naltrexone disrupted associations between cingulate perfusion and ketamine’s subjective or antidepressant effects without preventing the regional perfusion increases.^56^ This identifies an interaction between opioid function and cortical response that is distinct from the thalamic δ-opioid explanation for analgesia. The contribution of opioid pharmacology may therefore differ between pain relief and mood improvement.

Prefrontal adenosine A1 activity may modify arousal through its control of cholinergic transmission. ADORA1 was represented in the S-ketamine, R-ketamine, and hydroxyketamine fingerprints; no eligible ketamine–ADORA1 activity measurement was available for the parent-compound calculation. In mice, prefrontal A1-receptor blockade increased acetylcholine release and wakefulness and shortened recovery from isoflurane anesthesia, while A1 activation reduced acetylcholine release in the pontine reticular formation.^84^ Prefrontal adenosine can thus influence both local cortical activity and descending regulation of arousal. An A1-mediated reduction in acetylcholine would favor sedation, whereas reduced A1 activity would favor wakefulness. Comparing the direction and strength of A1 activity could therefore help explain arousal-related differences among ketamine enantiomers and metabolites.

Several limitations affect the interpretation of these findings. HR balances the published affinities of ketamine for its receptors with a numeric expression value in each tissue, but pharmacological measurements are incomplete and differ in species and assay conditions,^37^ so the absence of a feature can reflect missing measurements or the statistical threshold used to select the fingerprint. Tissue expression estimates do not specify protein abundance, receptor assembly, or the neuronal populations expressing a target.^92–94^ Importantly, HR does not distinguish directionality (agonism versus antagonism), only intensity of activity. For prefrontal NMDARs, inhibition of interneurons and pyramidal cells can have different consequences.^58^ HR scores also omit regional drug concentrations and the duration and direction of target engagement. Shared fingerprint components can therefore accompany different effects when compounds differ in exposure or in how they modify receptor activity.^95^ Cell-resolved expression, protein measurements, and functional pharmacology would help distinguish the cellular actions responsible for the proposed therapeutic and adverse effects. Once an interesting target-tissue pair is implicated in a component phenotype, further research is needed to determine directionality in the MoA.

Overall, this study provides a map of ketamine’s activity on specific receptors and signaling systems in specific brain regions. The map is complex but may be dissected to better understand or hypothesize about ketamine’s MoA against pain and depression, as well as its hallucinations or other phenotypes.

### Literature search

The Discussion was informed by a broad manual review of pharmacological, behavioral, and clinical literature. PubMed, Europe PMC, and PubMed Central were searched, with OpenAlex, Crossref, and publisher records used to identify and check sources. Search terms combined ketamine, its enantiomers or metabolites, receptor names, gene symbols, and anatomical regions with terms for depression, bipolar disorder, social behavior, pain and nociception, anesthesia, dissociation, hallucinations, memory and cognition, and arousal and sedation. Additional searches examined relevant receptor functions independently of ketamine exposure. Reference lists of relevant primary articles and reviews were examined for further studies. Evidence was considered according to the compound, receptor or receptor complex, species, anatomical region, and outcome examined.

## ABBREVIATIONS

5-HT: serotonin (5-hydroxytryptamine)
AC50: half-maximal activity concentration
AMPA: α-amino-3-hydroxy-5-methyl-4-isoxazolepropionic acid
AUD: alcohol use disorder
CNS: central nervous system
DOB: 2,5-dimethoxy-4-bromoamphetamine
EC50: half-maximal effective concentration
EM-SVD: expectation-maximization singular value decomposition
FDA: U.S. Food and Drug Administration
GABA: γ-aminobutyric acid
GCRMA: GC robust multi-array average
HR: historeceptomics
IC50: half-maximal inhibitory concentration
IUPHAR/BPS: International Union of Basic and Clinical Pharmacology/British Pharmacological Society
IV: intravenous
Kd: dissociation constant
Ki: inhibition constant
LSD: lysergic acid diethylamide
MDD: major depressive disorder
MoA: mechanism of action
mTOR: mechanistic target of rapamycin
NIMH: National Institute of Mental Health
NMDA: N-methyl-D-aspartate
NMDAR: NMDA receptor
OCD: obsessive-compulsive disorder
PCA: principal component analysis
PFC: prefrontal cortex
PTSD: post-traumatic stress disorder
RCT: randomized controlled trial
rs-fMRI: resting-state functional magnetic resonance imaging
SERT: serotonin transporter
TRD: treatment-resistant depression.

## Author contributions

**TC conceived the study. ER, JF, and HA performed the analyses. Funding: None declared**

## Acknowledgments

**None**

## Ethics approval

**Not applicable. This study did not involve human participants or animal experimentation**.

## Data and code availability

**Analysis code, documentation, test fixtures, and selected reference outputs are available in the Ketamine Historeceptomics repository: https://github.com/ericrosenn1/ketamine-historeceptomics**.

## AI use statement

**Generative AI tools (ChatGPT, GitHub Copilot) were used for language editing, improvement of manuscript readability, and GitHub repository organization and documentation. The scientific content, analyses, results, and conclusions were developed and verified by the authors**.

