## Supplement for "Historeceptomics Profiling of Ketamine"

#### Clinical dosing and effects

**Table S1.** Ketamine prescription dosing and effects.

| Use case | Normal dosing | Expected effects | Severe effects | Ref. |
|---|---|---|---|---|
| Acute analgesia | 0.1–0.3 mg/kg IV; 0.5–1 mg/kg intranasal | Analgesia with consciousness usually preserved; may reduce opioid requirements | Dizziness, nausea, perceptual effects, sedation, and transient BP/heart-rate elevation increase with dose and rapid administration | Morgan et al. (2021); Schwenk et al. (2018) |
| Chronic / neuropathic pain (off-label) | Protocol-dependent; often starts below 0.5 mg/kg IV, sometimes prolonged or repeated infusions | Reduction of refractory or neuropathic pain; possible opioid-sparing effect | Psychomimetic effects, somnolence, hypertension, nausea; repeated/prolonged exposure raises hepatic and urinary concerns | Cohen et al. (2018) |
| Antidepressant (off-label) | 0.5 mg/kg IV over ~40 min; repeated trials often twice or three times weekly | Rapid antidepressant response, often within hours; benefit may persist several days | Transient dissociation, dizziness, nausea, sedation, and BP elevation; higher doses generally produce more dissociation/hemodynamic effects | Berman et al. (2000); Zarate et al. (2006); Singh et al. (2016) |
| Procedural dissociation / deep sedation | ≈1–2 mg/kg IV or 3–4 mg/kg IM | Dissociation, profound analgesia, amnesia, reduced responsiveness | Salivation, vomiting, emergence reactions, hypertension, tachycardia; occasional airway or respiratory complications | Ketalar (2026); Cappellini et al. (2024) |
| General anesthesia | FDA label: 1–4.5 mg/kg IV; 6.5–13 mg/kg IM. Typical IV induction ≈2 mg/kg | Surgical anesthesia; ~2 mg/kg IV generally yields ~5–10 min of anesthesia | Higher doses prolong recovery; rapid IV delivery increases respiratory-depression and pressor-response risk | Ketalar (2026) |
| Nonmedical / recreational exposure | No standardized safe dose; 1 mg/kg IV overlaps the clinical dissociative-sedation range | Euphoria, dissociation, perceptual alteration, impaired coordination, amnesia | Risk rises with rapid delivery, unknown purity, and CNS-depressant combinations; repeated use is associated with dependence, cognitive, urinary, and hepatobiliary injury | Cappellini et al. (2024); Erskine et al. (2026) |

*Note.* IV, intravenous; IM, intramuscular; BP, blood pressure.

#### Supplementary data sources and analytical inputs

**Table S2.** Data sources and analytical incorporation in the historeceptomic workflow.

| Data component | Resource | Use in analysis | Primary citation / DOI | Numerically incorporated into final HR matrix? |
|---|---|---|---|---|
| ChEMBL | ChEMBL | Selected pharmacology and provenance | 10.1093/nar/gkad1004 | Yes; 1 selected row |
| PubChem BioAssay | PubChem BioAssay | Selected assay pharmacology and provenance | 10.1093/nar/gkae1059; 10.1093/nar/gkw1118 | Yes; 25 selected rows |
| NIMH PDSP Ki Database | NIMH Psychoactive Drug Screening Program Ki Database | Selected activity, species, endpoint, and relation records | 10.1177/107385840000600408 | Yes; 36 selected rows |
| IUPHAR/BPS | Guide to PHARMACOLOGY | Target and pharmacology source context | 10.1093/nar/gkaf1067 | No directly selected row |
| BindingDB | BindingDB | Audited candidate pharmacology | 10.1093/nar/gkae1075 | No |
| Primary literature | Publisher and repository records | Source-level pharmacology provenance | 10.1021/acs.jmedchem.0c01193; 10.1038/s41467-023-40064-9 | Yes; selected records with DOI provenance |
| Human expression | BioGPS GeneAtlas U133A/GNF1H | 77-tissue human expression panel | 10.1073/pnas.0400782101; 10.1186/gb-2009-10-11-r130; 10.1093/nar/gkv1104 | Yes |
| GTEx v8 | GTEx v8 | Supporting source-specific sensitivity resource | 10.1126/science.aaz1776 | No; not combined with final BioGPS matrix |
| Allen Human Brain Atlas | Allen Human Brain Atlas | Supporting brain-expression resource | 10.1038/nature11405 | No; not combined with final BioGPS matrix |
| Outlier procedure | Rosner generalized ESD | One-sided upper-tail fingerprint selection | 10.1080/00401706.1983.10487848 | No; applied after HR-score construction |

*Note.* Resources listed as not numerically incorporated were retained for source context or supporting analyses but were not combined into the final BioGPS-based HR matrix. HR, historeceptomic; ESD, extreme Studentized deviate; PDSP, Psychoactive Drug Screening Program; GTEx, Genotype-Tissue Expression.

**Table S3.** Ketamine-family analytical profiles and HR fingerprint call counts.

| Order | Analytical profile | Profile definition | Calls at α = 0.001 | Calls at α = 0.0001 |
|---|---|---|---|---|
| 1 | Ketamine (primary profile) | Source-confirmed racemic and stereochemistry-unspecified ketamine measurements | 19 | 14 |
| 2 | Ketamine (confirmed racemate) | Measurements explicitly identified as racemic ketamine | 12 | 10 |
| 3 | S-ketamine (esketamine) | S-enantiomer | 12 | 9 |
| 4 | R-ketamine (arketamine) | R-enantiomer | 11 | 9 |
| 5 | Hydroxyketamine, unspecified-isomer aggregate | Hydroxyketamine measurements not resolved to one isomer | 7 | 7 |
| 6 | (2R,6R)-hydroxynorketamine | Stereochemically resolved hydroxynorketamine | 0 | 0 |
| 7 | (2S,6S)-hydroxynorketamine | Stereochemically resolved hydroxynorketamine | 0 | 0 |
| 8 | Hydroxynorketamine, generic identity | Generic HNK identity | 0 | 0 |
| 9 | Hydroxyketamine, generic identity | Generic hydroxyketamine identity | 0 | 0 |
| 10 | Norketamine | N-demethylated ketamine metabolite | 1 | 1 |

*Note.* Fingerprint calls were selected using one-sided upper-tail generalized ESD. The primary threshold was α = 0.001; α = 0.0001 was the stricter sensitivity threshold.

#### Ketamine CNS and whole-body fingerprint data

**Table S4.** Ketamine strict-CNS historeceptomic fingerprint and GESD statistics.

| Rank | Target | Tissue | HR score | Expression z | GESD R | Critical λ | R - λ | Retained at α = 0.0001 |
|---|---|---|---|---|---|---|---|---|
| 1 | GRIN1 | Prefrontal cortex | 51.883 | 7.692 | 10.392 | 4.737 | 5.654 | Yes |
| 2 | CHRM3 | Prefrontal cortex | 42.331 | 8.466 | 8.964 | 4.737 | 4.227 | Yes |
| 3 | GRIN2A | Subthalamic nucleus | 41.158 | 6.102 | 9.079 | 4.737 | 4.342 | Yes |
| 4 | HTR2C | Caudate nucleus | 35.091 | 7.018 | 8.078 | 4.737 | 3.341 | Yes |
| 5 | HTR2A | Prefrontal cortex | 34.018 | 6.804 | 8.094 | 4.737 | 3.358 | Yes |
| 6 | GABRA2 | Amygdala | 30.794 | 6.809 | 7.580 | 4.736 | 2.844 | Yes |
| 7 | OPRD1 | Thalamus | 23.883 | 4.777 | 6.069 | 4.736 | 1.333 | Yes |
| 8 | GRIN1 | Amygdala | 23.708 | 3.515 | 6.138 | 4.736 | 1.402 | Yes |
| 9 | SLC6A3 | Hypothalamus | 21.896 | 4.841 | 5.785 | 4.736 | 1.049 | Yes |
| 10 | DRD1 | Caudate nucleus | 20.442 | 4.088 | 5.499 | 4.736 | 0.764 | Yes |
| 11 | GRIN2C | Cerebellar peduncles | 20.073 | 2.976 | 5.486 | 4.735 | 0.750 | Yes |
| 12 | DRD1 | Pons | 19.217 | 3.843 | 5.337 | 4.735 | 0.602 | Yes |
| 13 | GABRA1 | Occipital lobe | 19.070 | 4.216 | 5.375 | 4.735 | 0.640 | Yes |
| 14 | GRIN2C | Amygdala | 18.453 | 2.736 | 5.282 | 4.735 | 0.547 | Yes |
| 15 | GRIN2A | Parietal lobe | 17.328 | 2.569 | 5.039 | 4.734 | 0.305 | No |
| 16 | PDE4A | Prefrontal cortex | 16.197 | 3.581 | 4.781 | 4.734 | 0.046 | No |
| 17 | GABRA1 | Parietal lobe | 15.825 | 3.499 | 4.729 | 4.734 | -0.005 | No |
| 18 | HTR2C | Hypothalamus | 15.665 | 3.133 | 4.737 | 4.734 | 0.004 | No |
| 19 | GRIN2A | Cingulate cortex | 15.477 | 2.295 | 4.737 | 4.734 | 0.003 | No |

*Note.* All rows are target-tissue fingerprint calls selected at α = 0.001. HR score is pActivity multiplied by the standardized tissue-expression z score. GESD R is the observed upper-tail test statistic, λ is the corresponding critical value, and R - λ is the decision margin.

**Table S5.** Ketamine whole-body historeceptomic fingerprint.

| Rank | Target | Tissue | Tissue class | HR score | Retained at α = 0.0001 |
|---|---|---|---|---|---|
| 1 | GRIN1 | Prefrontal cortex | CNS | 51.88 | Yes |
| 2 | GRIN2C | Thyroid | Peripheral/non-CNS | 44.13 | Yes |
| 3 | CHRM3 | Prefrontal cortex | CNS | 42.33 | Yes |
| 4 | GRIN2A | Subthalamic nucleus | CNS | 41.16 | Yes |
| 5 | F2 | Liver | Peripheral/non-CNS | 39.10 | Yes |
| 6 | SLC6A4 | Pineal gland, nighttime | Peripheral/non-CNS | 35.83 | Yes |
| 7 | NR1I2 | Liver | Peripheral/non-CNS | 35.36 | Yes |
| 8 | HTR1A | Liver | Peripheral/non-CNS | 35.25 | Yes |
| 9 | PTGS1 | Smooth muscle | Peripheral/non-CNS | 35.21 | Yes |
| 10 | HTR2C | Caudate nucleus | CNS | 35.09 | Yes |
| 11 | AR | Liver | Peripheral/non-CNS | 35.05 | Yes |
| 12 | ADRB2 | CD56+ natural killer cells | Peripheral/non-CNS | 34.60 | Yes |
| 13 | HTR2A | Prefrontal cortex | CNS | 34.02 | Yes |
| 14 | BCHE | Fetal liver | Peripheral/non-CNS | 32.78 | Yes |
| 15 | ADRB1 | Pineal gland, nighttime | Peripheral/non-CNS | 32.55 | Yes |
| 16 | GABRA2 | Amygdala | CNS | 30.79 | Yes |
| 17 | GABRG2 | Pineal gland, daytime | Peripheral/non-CNS | 30.03 | Yes |
| 18 | OPRM1 | Superior cervical ganglion | Peripheral/non-CNS | 29.80 | Yes |
| 19 | GRIN3A | Uterus | Peripheral/non-CNS | 29.22 | Yes |

| Rank | Target | Tissue | Tissue class | HR score | Retained at α = 0.0001 |
|---|---|---|---|---|---|
| 20 | HTR3A | Dorsal root ganglion | Peripheral/non-CNS | 28.20 | Yes |
| 21 | CHRM2 | Skeletal muscle | Peripheral/non-CNS | 27.16 | Yes |
| 22 | GRIN3A | Skeletal muscle | Peripheral/non-CNS | 26.34 | Yes |
| 23 | ABCC3 | Adrenal cortex | Peripheral/non-CNS | 26.30 | Yes |
| 24 | BCHE | Uterus | Peripheral/non-CNS | 26.09 | Yes |
| 25 | HRH2 | Cardiac myocytes | Peripheral/non-CNS | 24.97 | Yes |
| 26 | GABRB2 | Liver | Peripheral/non-CNS | 24.67 | Yes |
| 27 | ADORA3 | Testis interstitial cells | Peripheral/non-CNS | 24.59 | Yes |
| 28 | HRH2 | Heart | Peripheral/non-CNS | 24.26 | Yes |
| 29 | ABCC2 | Pancreatic islets | Peripheral/non-CNS | 24.10 | Yes |
| 30 | GABRG2 | Pineal gland, nighttime | Peripheral/non-CNS | 24.05 | Yes |
| 31 | OPRD1 | Thalamus | CNS | 23.88 | Yes |
| 32 | ABCC4 | CD71+ early erythroid cells | Peripheral/non-CNS | 23.85 | Yes |
| 33 | HRH1 | Superior cervical ganglion | Peripheral/non-CNS | 23.78 | Yes |
| 34 | GRIN1 | Amygdala | CNS | 23.71 | Yes |
| 35 | MAOA | Placenta | Peripheral/non-CNS | 23.69 | Yes |
| 36 | HTR2B | Uterine corpus | Peripheral/non-CNS | 22.28 | Yes |
| 37 | ADRA2A | Fetal brain | Peripheral/non-CNS | 22.27 | Yes |
| 38 | KDR | Thyroid | Peripheral/non-CNS | 22.01 | Yes |
| 39 | SLC6A3 | Hypothalamus | CNS | 21.90 | No |
| 40 | ADRA2A | Lymph node | Peripheral/non-CNS | 21.67 | No |
| 41 | GRIN3A | Fetal brain | Peripheral/non-CNS | 21.54 | No |
| 42 | MAOA | Small intestine | Peripheral/non-CNS | 21.42 | No |
| 43 | ADRA1A | Cardiac myocytes | Peripheral/non-CNS | 21.35 | No |
| 44 | ABCC4 | Prostate | Peripheral/non-CNS | 21.30 | No |
| 45 | GRIN2B | Skeletal muscle | Peripheral/non-CNS | 21.29 | No |
| 46 | ACHE | Pineal gland, nighttime | Peripheral/non-CNS | 21.22 | No |
| 47 | DRD1 | Caudate nucleus | CNS | 20.44 | No |
| 48 | DRD3 | Heart | Peripheral/non-CNS | 20.39 | No |
| 49 | KDR | Retina | Peripheral/non-CNS | 20.22 | No |
| 50 | ADRB1 | Pineal gland, daytime | Peripheral/non-CNS | 20.10 | No |
| 51 | ADRA2C | Skeletal muscle | Peripheral/non-CNS | 20.10 | No |
| 52 | GRIN2C | Cerebellar peduncles | CNS | 20.07 | No |
| 53 | PDE4A | Pineal gland, nighttime | Peripheral/non-CNS | 19.94 | No |
| 54 | ADORA3 | Testis germ cells | Peripheral/non-CNS | 19.90 | No |
| 55 | HTR2B | Uterus | Peripheral/non-CNS | 19.82 | No |
| 56 | ACHE | Adipocytes | Peripheral/non-CNS | 19.59 | No |
| 57 | KCNH2 | CD34+ cells | Peripheral/non-CNS | 19.39 | No |
| 58 | DRD1 | Pons | CNS | 19.22 | No |
| 59 | GABRA1 | Occipital lobe | CNS | 19.07 | No |

*Note.* All 59 rows were selected independently from the 58-target by 77-tissue whole-body HR-score matrix at α = 0.001. The sensitivity column identifies the 38 calls also retained at α = 0.0001.

### Ketamine-family fingerprint comparisons

**Table S6.** Selected fingerprint overlap within the ketamine family.

This table summarizes selected pairwise fingerprint comparisons within the ketamine family using call counts, Jaccard similarity, and the overlap coefficient. These values provide the quantitative basis for the family-comparison results and interpretation of shared and compound-specific fingerprint features.

| Comparison | Calls A | Calls B | Shared calls | Union | Jaccard | Overlap coefficient |
|---|---|---|---|---|---|---|
| Ketamine (primary profile) vs Ketamine (confirmed racemate) | 19 | 12 | 10 | 21 | 0.48 | 0.83 |
| Ketamine (primary profile) vs S-ketamine (esketamine) | 19 | 12 | 10 | 21 | 0.48 | 0.83 |
| Ketamine (primary profile) vs R-ketamine (arketamine) | 19 | 11 | 9 | 21 | 0.43 | 0.82 |
| S-ketamine (esketamine) vs R-ketamine (arketamine) | 12 | 11 | 11 | 12 | 0.92 | 1.00 |
| Ketamine (primary profile) vs Hydroxyketamine, unspecified-isomer aggregate | 19 | 7 | 5 | 21 | 0.24 | 0.71 |
| Ketamine (confirmed racemate) vs S-ketamine (esketamine) | 12 | 12 | 8 | 16 | 0.50 | 0.67 |
| Ketamine (confirmed racemate) vs R-ketamine (arketamine) | 12 | 11 | 7 | 16 | 0.44 | 0.64 |
| S-ketamine (esketamine) vs Hydroxyketamine, unspecified-isomer aggregate | 12 | 7 | 7 | 12 | 0.58 | 1.00 |
| R-ketamine (arketamine) vs Hydroxyketamine, unspecified-isomer aggregate | 11 | 7 | 7 | 11 | 0.64 | 1.00 |
| Ketamine (primary profile) vs Norketamine | 19 | 1 | 1 | 19 | 0.05 | 1.00 |

*Note.* Values use α = 0.001 fingerprint calls. Jaccard similarity equals shared calls divided by the call-set union. The overlap coefficient equals shared calls divided by the size of the smaller nonempty fingerprint.

**A** **B**

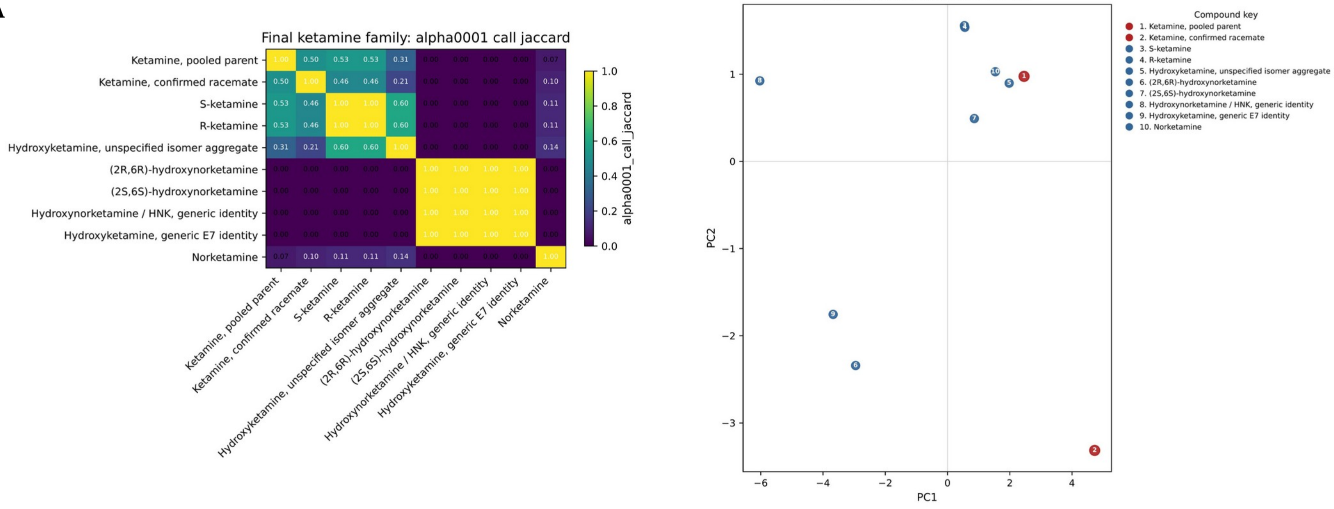


**Figure S1. Ketamine-family fingerprint sensitivity analyses.** (A) Call-set Jaccard similarity and (B) sparse fingerprint PCA at α = 0.0001 for the 10 ketamine-family profiles. These analyses assess whether the principal family-level similarity pattern and ordination remain evident under the stricter fingerprint threshold.

#### External-drug fingerprint comparisons

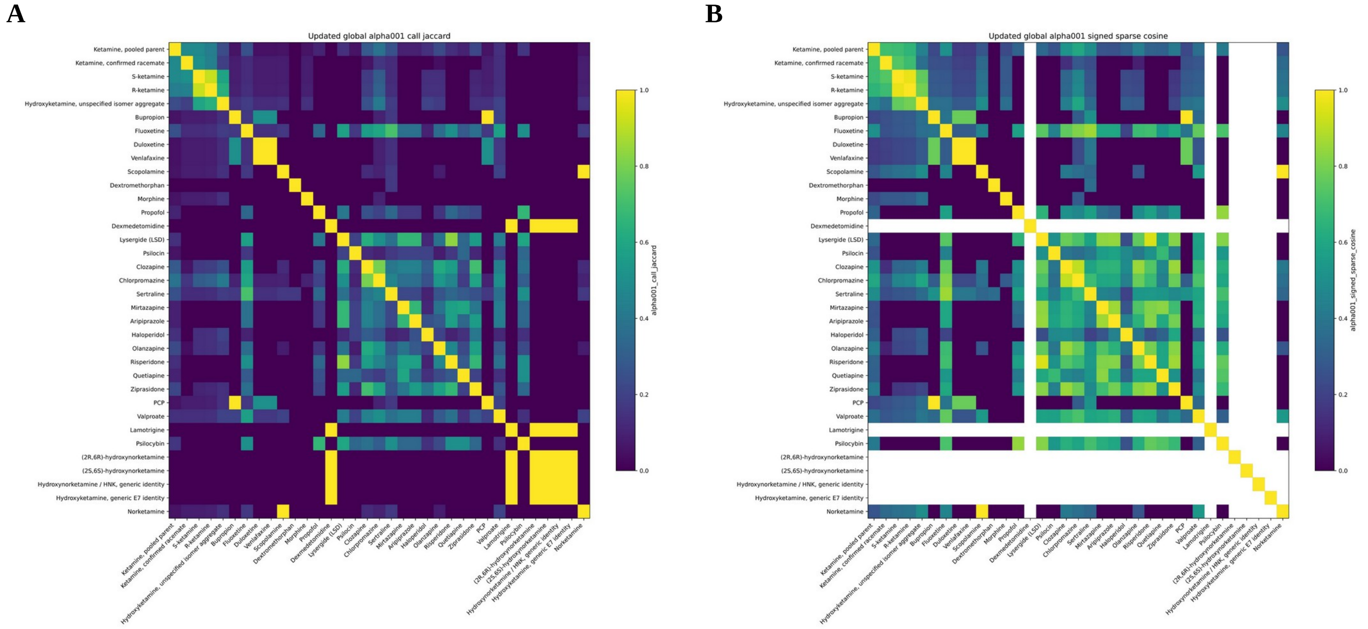


**Figure S2. Primary external-drug fingerprint comparisons.** (A) Call-set Jaccard similarity and (B) signed sparse-cosine similarity at α = 0.001 across the final 35-profile combined roster. These comparisons were used to position ketamine-family fingerprints relative to established psychoactive reference drugs and support the external-drug analyses in the manuscript.

**Table S7.** Fingerprint overlap between ketamine and psychoactive reference drugs.

This table reports α = 0.001 fingerprint overlap between the primary ketamine profile and all 25 psychoactive reference drugs. It was used to identify the strongest reference-drug overlaps and to support the comparative pharmacology analysis in the manuscript.

| Reference drug | Drug group | Reference calls | Shared calls | Union | Jaccard | Overlap coefficient |
|---|---|---|---|---|---|---|
| Bupropion | Antidepressant | 1 | 1 | 19 | 0.05 | 1.00 |
| Fluoxetine | Antidepressant | 6 | 5 | 20 | 0.25 | 0.83 |
| Duloxetine | Antidepressant | 2 | 1 | 20 | 0.05 | 0.50 |
| Venlafaxine | Antidepressant | 2 | 1 | 20 | 0.05 | 0.50 |
| Scopolamine | Cholinergic | 1 | 1 | 19 | 0.05 | 1.00 |
| Dextromethorphan | NMDA/dissociative | 1 | 0 | 20 | 0.00 | 0.00 |
| Morphine | Analgesic/opioid | 1 | 1 | 19 | 0.05 | 1.00 |
| Propofol | Anesthetic/sedative | 2 | 2 | 19 | 0.11 | 1.00 |
| Dexmedetomidine | Anesthetic/sedative | 0 | 0 | 19 | 0.00 | 0.00 |
| Lysergide (LSD) | Serotonergic psychedelic | 5 | 3 | 21 | 0.14 | 0.60 |
| Psilocin | Serotonergic psychedelic | 1 | 1 | 19 | 0.05 | 1.00 |
| Clozapine | Antipsychotic | 8 | 6 | 21 | 0.29 | 0.75 |
| Chlorpromazine | Antipsychotic | 10 | 8 | 21 | 0.38 | 0.80 |
| Sertraline | Antidepressant | 6 | 5 | 20 | 0.25 | 0.83 |
| Mirtazapine | Antidepressant | 5 | 2 | 22 | 0.09 | 0.40 |
| Aripiprazole | Antipsychotic | 5 | 2 | 22 | 0.09 | 0.40 |
| Haloperidol | Antipsychotic | 5 | 3 | 21 | 0.14 | 0.60 |
| Olanzapine | Antipsychotic | 13 | 6 | 26 | 0.23 | 0.46 |
| Risperidone | Antipsychotic | 6 | 3 | 22 | 0.14 | 0.50 |
| Quetiapine | Antipsychotic | 3 | 2 | 20 | 0.10 | 0.67 |
| Ziprasidone | Antipsychotic | 9 | 5 | 23 | 0.22 | 0.56 |
| PCP | NMDA/dissociative | 1 | 1 | 19 | 0.05 | 1.00 |
| Valproate | Mood stabilizer/antiepileptic | 5 | 3 | 21 | 0.14 | 0.60 |
| Lamotrigine | Mood stabilizer/antiepileptic | 0 | 0 | 19 | 0.00 | 0.00 |
| Psilocybin | Serotonergic psychedelic | 3 | 3 | 19 | 0.16 | 1.00 |

*Note.* All comparisons use the 19-call ketamine primary fingerprint and the final 35-profile α = 0.001 analysis. Drug groups organize the reference set and do not assign ketamine to a pharmacological class.

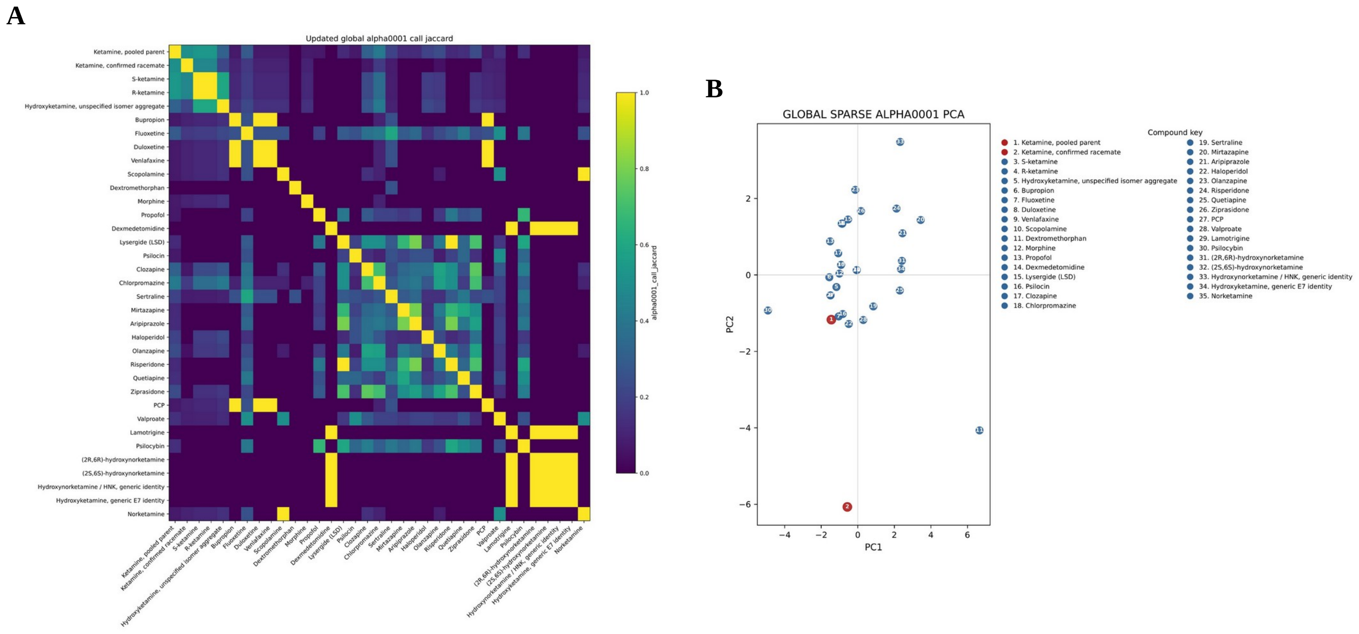


**Figure S3. External-drug fingerprint sensitivity analyses.** (A) Call-set Jaccard similarity and (B) sparse fingerprint PCA at α = 0.0001 across the final 35-profile combined roster. These sensitivity analyses test the stability of external-drug fingerprint similarity and multivariate positioning at the stricter threshold.

### CNS phenotype mapping

**Table S8.** CNS phenotype mapping summary.

This table summarizes the design and yield of the CNS phenotype-mapping analysis, from the 400 predefined combinations to the accepted mappings and Sankey paths. It provides the denominators and retained-path counts used to interpret the phenotype-mapping figures.

| Measure | Count |
|---|---|
| Fingerprint target-tissue coordinates evaluated | 20 |
| CNS effects evaluated | 20 |
| Combinations searched | 400 |
| Accepted target-tissue-phenotype mappings | 32 |
| Target-tissue pairs with accepted mappings | 12 |
| Exact phenotype categories | 15 |
| Compact phenotype nodes | 12 |
| Full compound-pair-phenotype paths | 76 |
| Compact compound-pair-phenotype paths | 71 |

*Note.* Only accepted target-tissue-phenotype mappings are represented in the Sankey figures. Path counts do not encode effect magnitude or evidence strength.
